# Identifying compositional and density changes across the Murine and Human Dentin-enamel Junction

**DOI:** 10.1101/2024.01.30.578062

**Authors:** Michael Truhlar, Bradley Rosenberg, Sobhan Katebifar, Roland Kroger, Alix Deymier

## Abstract

Human and mouse incisors are both primarily composed of dentin and enamel which meet at an interface called the dentin-enamel junction (DEJ). However, incisors in the two species have very different growth patterns, structures, and loading requirements. Since the DEJ is responsible for minimizing cracking at this at-risk interface between mechanically dissimilar dentin and enamel, its structure is expected to be significantly different between humans and mice. Here, strucutral and compositional gradients across human and murine incisors DEJs were measured via microcomputed tomography and Raman spectroscopy. Density gradients across the DEJ were significantly larger in humans compared to murine teeth likely due to the larger size of the mantle dentin. Multiple gradients in mineral content and crystallinity were found at the murine DEJ while the human DEJ only exhibited gradients in mineral content. Models predicting the modulus across the DEJ according to compositional results show that mineral crystallinity is critical in regulating the mechanical gradient across the murine DEJ. Together these results show the multiple ways in which the DEJ can adapt to variations in loading environment.

## Introduction

Incisors in most mammalian species are composed primarily of two phases: dentin and enamel. Dentin is a porous material composed of 70% mineral, 20% organic material (primarily collagen) and 10% fluids by weight[1]. Enamel is a dense tissue containing ∽96% mineral, 1% organic matrix, and 3% water by weight. In addition to the compositional differences the two differ greatly in their structures resulting in two mechanically dissimilar materials. Despite their differences, the two tissues contact each other via an interface termed the dentin-enamel junction (DEJ). Interfaces between mechanically dissimilar materials are prone to the formation of stress concentrations and associated failure. However, the DEJ appears to be highly resistant to fracture [2]. This seems to be due to presence of compositional, structural, and mechanical gradients across the interface [3-10]. The challenge has been accurately measuring changes in these DEJ gradients.

Establishing the width of the DEJ has been a challenge in the current literature. The results of previous studies have varied greatly based on their methodology. Functional stress testing through crack dissipation and mechanical stiffness testing have yielded results supposing a functional width in the range of 100 µm [3, 4]. Studies utilizing nanoindentation techniques to quantify the elastic modulus and hardness across the DEJ have consistently found a resulting DEJ size of 10-20 µm [5, 6] while those using micro-indentation have reported values of closer to the 100 µm [7]. SEM images have exposed the scalloped, micro-scalloped, and overlapping structure of the human DEJ. The overlapping nature of the DEJ and the micro-scallops indicates a variation of composition across the DEJ at a scale of 1-3 um [8-10]. Nano-scratch and x-ray spectroscopy techniques have been able to identify a gradient that exists at that approximate scale, with values around 1-2 um [8, 9]. It is clear from the varying reports of DEJ width, that it is highly dependent upon methodology and the manner in which the DEJ is defined.

This complexity is further compounded by the differences found in the DEJs of varying species. Studies have shown that composition and structure of the DEJ varies from one species to the next and also in evolutionary time [11-14]. This is especially important to consider when animals models are used as stand ins for the study of disease in human dental tissues. Mice are widely utilized as disease models thanks to the ease with which it is possible to modify their genome and the rapidity of breeding. However, in terms of dentition mice exhibit very different structures and growth patterns compared to humans. Human incisors, much like human molars, are single growth teeth composed of a dentin core coved by a crown of enamel. Mouse incisors are continuously growing with enamel present only on the labile surface and dentin on the lingual surface. These very different incisors are still used for the purpose of cutting food and other materials; however, in mice these teeth are made to wear down during feeding and gnawing behaviors. These significant differences are expected to lead to strong differences in the structure and mechanics of the DEJ. Thus, in this study we use a multimodal approach to investigate changes in gradients across the DEJ of murine and human teeth and relate them to the tissue mechanics.

## Methods

### Sample extraction and preparation

All animal protocols were completed in accordance with the UConn Health Institutional Animal Care and Use Committee. Lower incisors were surgically dissected from 4-5 month old CD-1 mice. Following extraction, the teeth were cleaned of any soft tissue and wrapped in gauze soaked in a phosphate buffered saline solution for storage at -4°C. The lower left incisors were then thawed and mounted onto glass slides utilizing Loctite superglue. The teeth were oriented on the slides such that the DEJ was perpendicular to the surface of the slide. Each sample was polished prior to Raman spectroscopy utilizing silicon carbide sandpaper, starting at grit 2,000 and increasing incrementally to 10,000. Following this, the samples were polished utilizing a Komet USA 94013F HP170 diamond embedded polishing wheel in a rotary tool set to 6,000 rpm. The right teeth were wrapped in Phosphate buffered saline (PBS) soaked gauze and frozen for microcomputed tomography (µCT).

Two human incisors were collected from the Education Support Services Anatomy Laboratory at the University of Connecticut School of Dental Medicine. The incisors were imaged via µCT and embedded in EpoKwick FC and polished using diamond slurries up to a grain size of 0.1 µm for Raman analysis.

### Microcomputed tomography

Murine teeth were thawed and imaged whole using a Scanco 50 µCT system. For the murine samples, a region near the middle of the tooth measuring 7.3 µm long was imaged at a resolution of 3.4 µm, Fig 1. For the human incisors, the crowns of both teeth were scanned with a resolution of 5 µm. The cross-sectional images of the murine and human teeth were examined using CTAn program (Bruker). For the murine samples, 5 sections were selected at 3 locations along the tooth: near the dental follicle (bottom), near the middle of the tooth (middle), and towards the exposed tip of the tooth (top). For each of these sections, 5 lines were drawn perpendicular to the DEJ starting from the pulp cavity and extending across the interface eventually reaching the edge of the enamel boundary, Fig 1. For the human teeth, 6 lines were collected across the DEJ in the mid-crown. Absorption intensity data as a function of position was then collected for each line. The data was truncated to maintain only gradients present between the enamel and dentin and analyzed using custom software in Matlab. This customizable code calculated the width of the gradient in absorption intensity data across the DEJ by fitting a logistic function to the intensity data and determining the full width half max (FWHM) of the derivative of that function.

**Figure 1:**
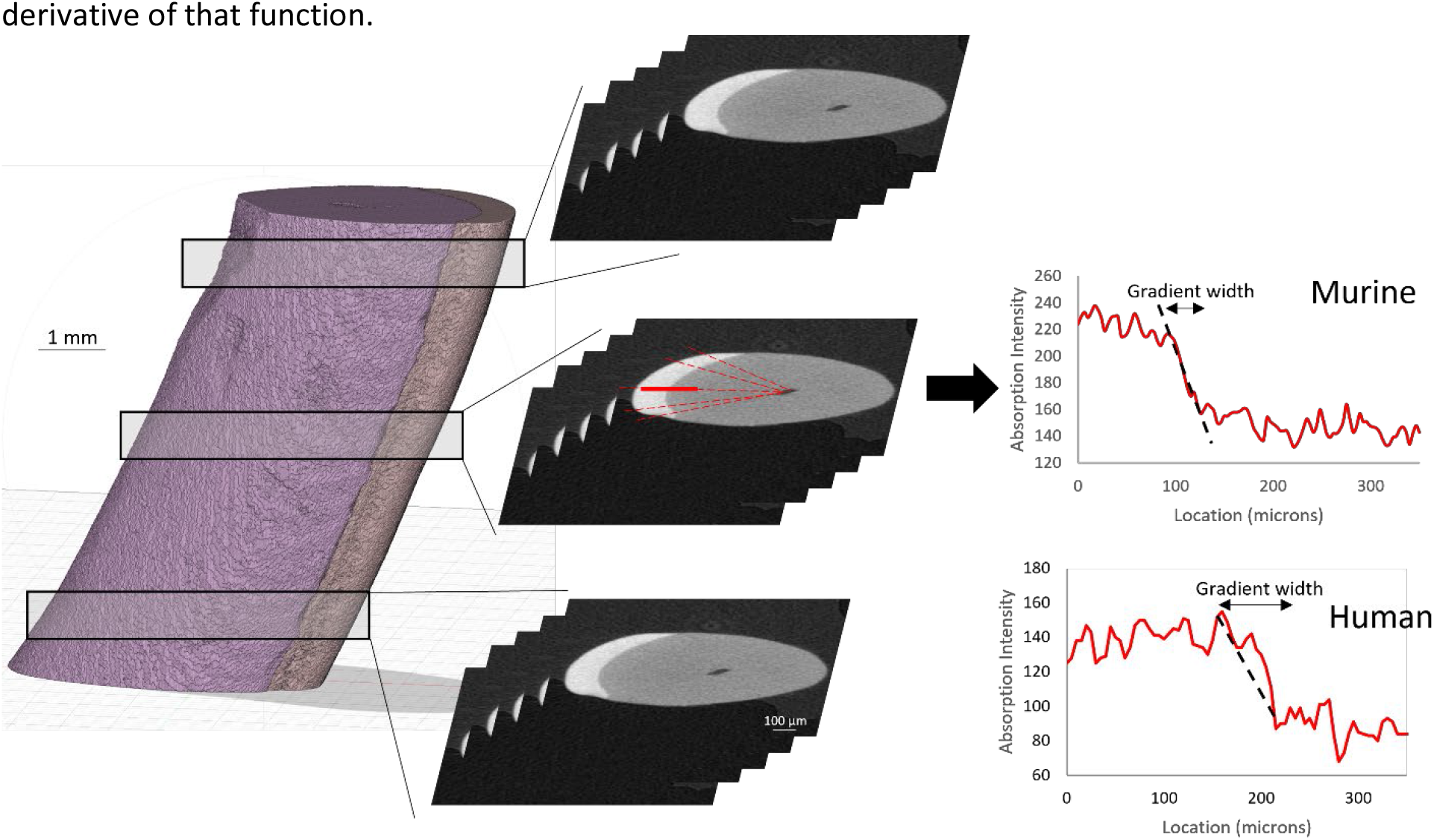
Schematic of the protocol for determining density gradients from murine teeth. uCT sections are selected from 3 regions in the tooth at which 25 lines are drawn across the DEJ. Representative plots of the absorption intensity across the DEJ is shown for murine and human samples.

### Raman Scans

Raman analysis was performed using a Witec Alpha 300 microscope paired with a Witec UHTS 400 spectrometer and a 785 nm laser. The white light view was used to identify the location of the DEJ in all samples, Fig 2. Once the approximate DEJ location was identified, we collected a 25×25 µm^2^ area scan with a resolution of 0.5 μm with an integration time of 20 sec for each sample. For the murine samples, one area scan was collected at the mid-point of the incisor length. For the human incisors, a 25 × 25 µm^2^ area scan was collected at the midheight of the crown on the labial side.

**Figure 2:**
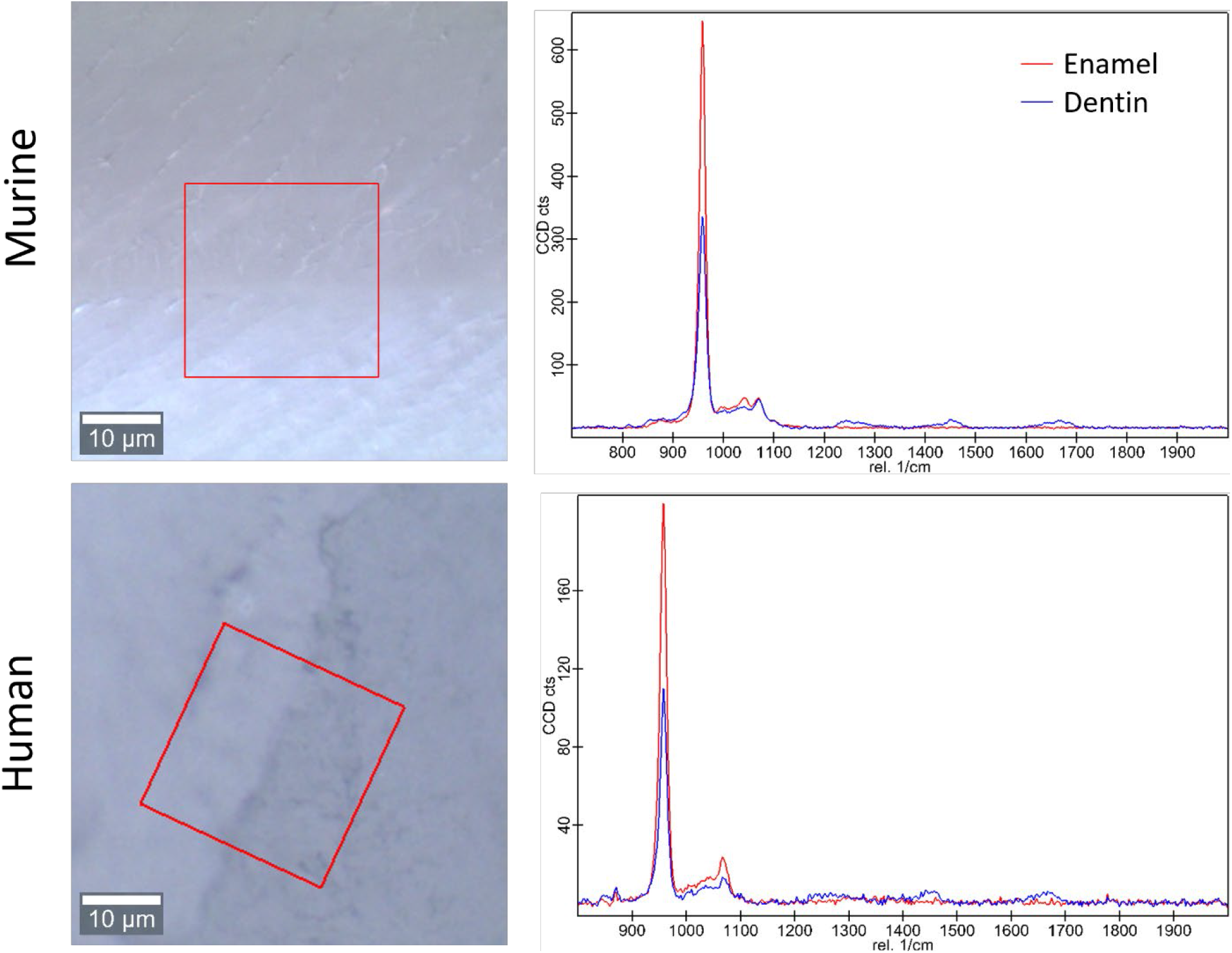
Bright field images of the DEJ in murine and human samples (left) showing the region of interest scanned for Raman spectroscopy. Representative Raman spectra for enamel and dentin are shown to the right.

### Area Scan Statistical Analysis

Following acquisition of these spectra, cosmic ray correction and background subtraction for the area scan data was done using Witec Program 5.1. The 960 Δcm^-1^ phosphate peak was fit as a Lorentzian peak to obtain the peak max intensity, width, and position. Custom Matlab code was then utilized to analyze these data. In summary, an experienced user was asked to provide a line perpendicular to the DEJ based on a 960 Δcm^-1^ phosphate peak intensity map. The angular orientation of this line was used to adjust the matrix creating plots of peak intensity, FWHM, and peak position as a function of location. The max intensity, peak width, and peak position as a function of location across the DEJ were then fit for all 50 lines utilizing a logistic regression. The derivative of this logistic fit was taken for each of our measurements to determine the center location of gradients (location of the max derivative value) as well as the width of the gradients of interest (full width half maximum value for each the derivative function), Fig 3. Values were averaged across al 50 lines for each sample.

**Figure 3:**
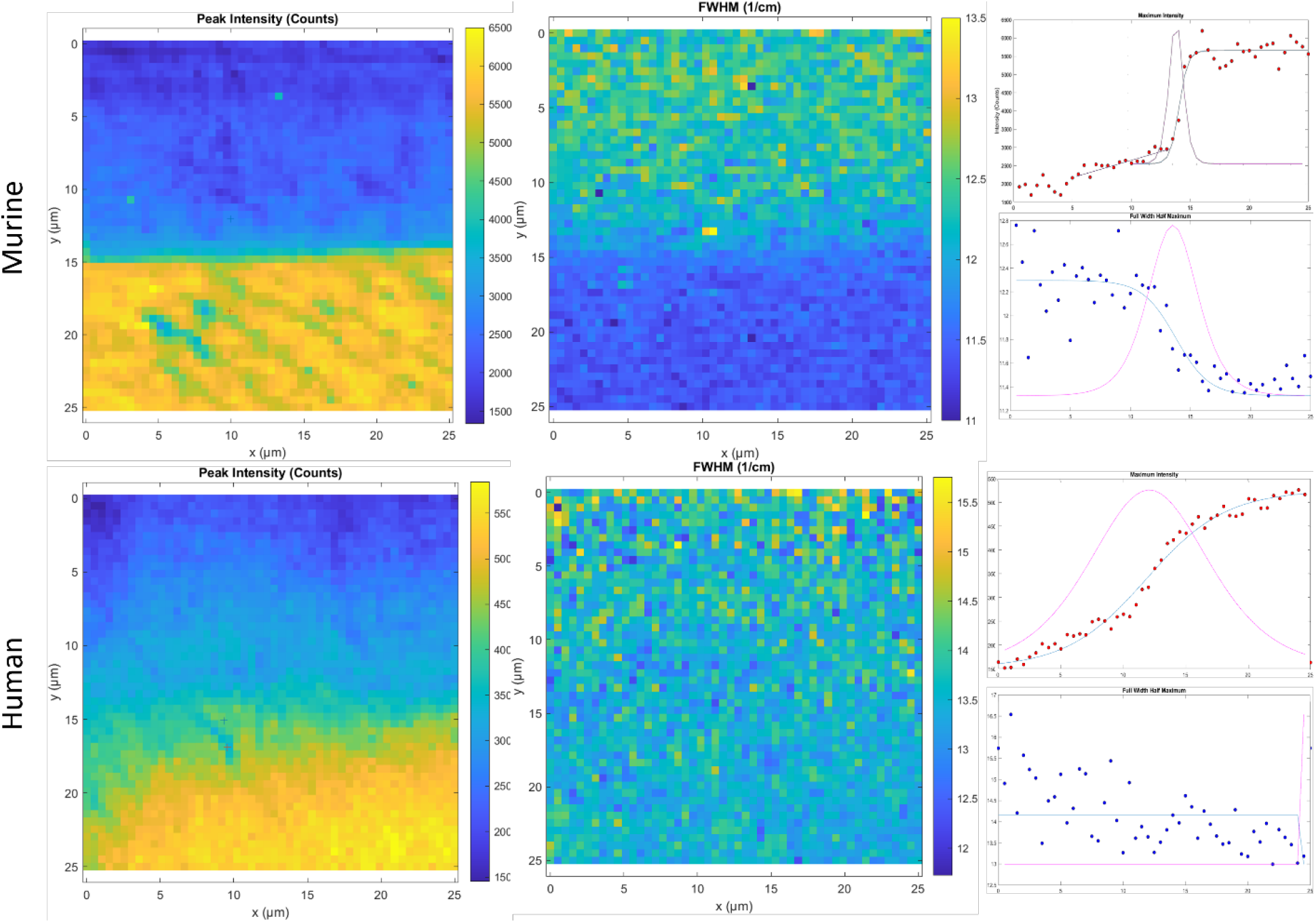
Representative maps of 960 peak intensity (left) and 960 peak FWHM (middle) showing the DEJ in mouse and human samples. To the right is shown the 960 peak intensity (red) and 960 peak FWHM (blue) fit for a single line across the DEJ for human and murine samples. Solid lines represent the logistic fit while the solid pink line is the derivative of the fit used to identify the gradient center and FWHM.

### Tissue Modulus

To understand the contribution of changes in mineral content and crystallinity on the DEJ mechanics, we developed a simplified composite model of dental tissues. To obtain information about the relative change in mineral content across the width of the DEJ, it was assumed that mineral content was proportional to the 960 peak maximum intensity. Therefore, mineral content as a function of position could be estimated from the average logistic fit of the 960 max intensity data. For the murine samples, only the 2^nd^ mineral content gradient was considered, and measures were made across 10 µm. For the human samples the single mineral content gradient was considered over 25 µm. Bounds were placed on this data such that pure dentin had a mineral content of 50% and pure enamel a mineral content of 100% [1]. The modulus of the tissue was then calculated as a function of location assuming that the material is being loaded in isostrain. Although highly simplistic, this shows the maximum effect of changes in mineral content and modulus to provide an upper bound. A single representative sample was used for each sample type.

To account for the effects of mineral crystallinity on the tissue mechanics, it was established that the mineral crystallinity is proportional to the fwhm of the 960 peak. Using previous data examining the effect of crystallinity on modulus [15, 16], a linear correlation was established between the width of the 960 peak and the mineral modulus. The logistic fit of the 960 FWHM data was used as a measure of the change in FWHM as a function of position across the DEJ. Together this data was used to calculate the mineral modulus as a function of position. For the murine samples which exhibit gradients in both 960 FWHM and max intensity, these values were then integrated into the isostrain calculations described above to determine the modulus of the tissue as a function of position, while accounting for contributions from the mineral crystallinity. From this we predicted the change in modulus based on crystallinity as a function of position across the DEJ. Using a span of 10 μm, the total tissue modulus was estimated by averaging the varying moduli across the tissue.

### Statistical Analysis

Gradient widths are reported as mean values ± the standard deviations. Comparisons between gradient width were performed using ANOVA followed by means comparison testing using the Tukey test.

Significance was set at p<0.05.

## Results

### Density Gradients from μCT

For the murine teeth, gradients in the tissue density as determined from μCT attenuation coefficients were measured across the DEJ at the top, bottom, and middle of the measured teeth. The average gradient widths were 24.4±0.72 μm for the bottom, 22.89±2.44 μm for the middle, and 22.33±3.34 μm for the top of the tooth, Fig 4A. However, none of these values were significantly different as a function of location across the tooth. Therefore, the total average value of the density gradient can be measured as 23.12±2.48 μm, Fig 4B.

**Figure 4:**
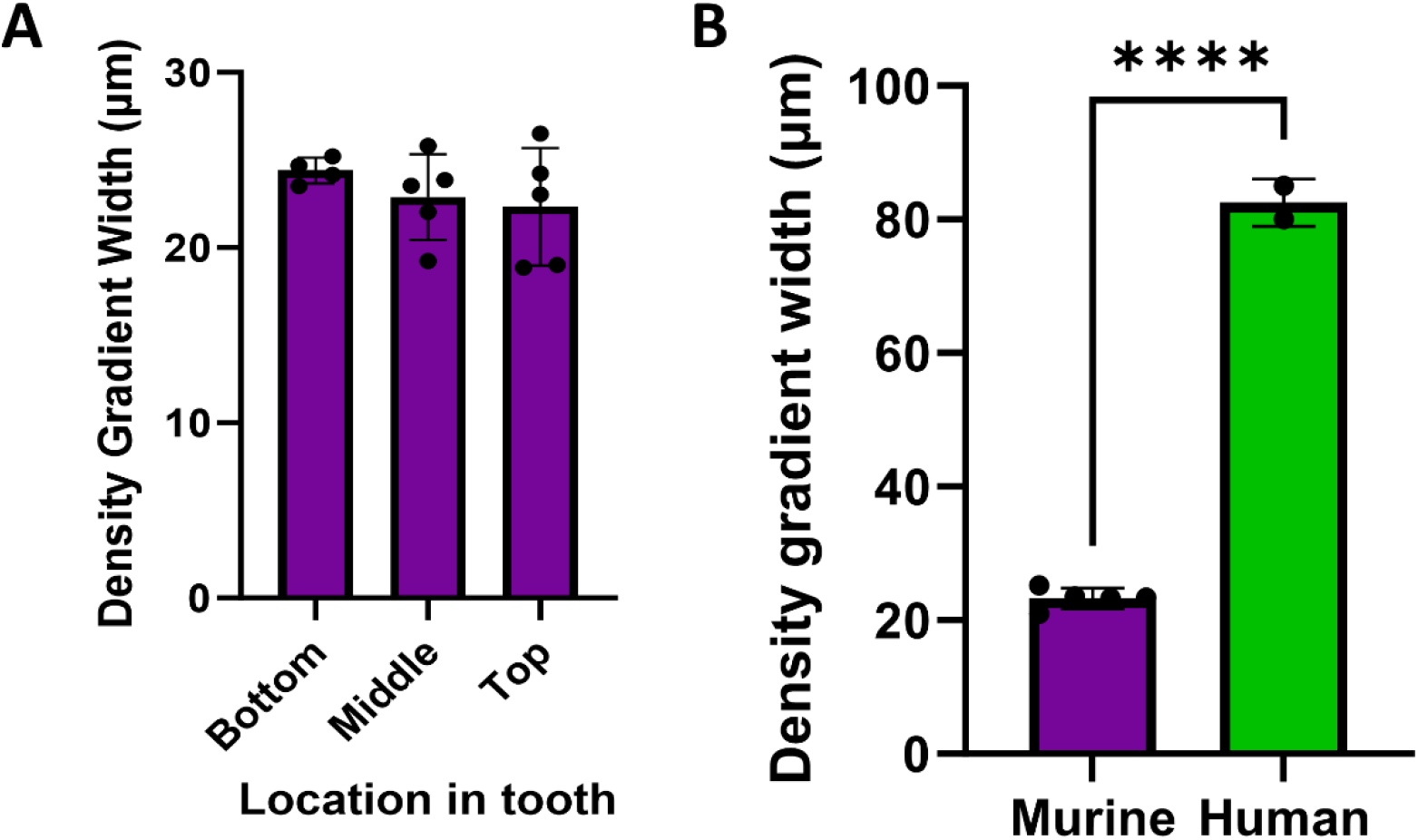
Plot (A) showing the width of the density gradient as measured from uCT at the three locations in the murine tooth. (B) Plot comparing the average murine desnity gradient width to the human average.

For the human incisors, the density gradient was measured at a single location in the mid-crown. The average density gradient width for the human incisors was 82.5±3.5 µm, Fig 4B.

### Phosphate Peak and FWHM Gradients

The gradient widths for both the phosphate peak intensity and phosphate peak FWHM were calculated for each sample. These two measurements are directly correlated to phosphate content (peak intensity) and mineral crystallinity (inverse of the FWHM). Upon visual inspection of the murine area scans it was seen that two distinct 960 Δcm^-1^ peak intensity gradients existed across the murine DEJ: an initial shallow gradient (1^st^ Max Int), followed by a second steeper gradient (2^nd^ Max Int), Fig 5. As a result, it became necessary to fit each of these gradients independently. The 1^st^ Max Int gradient was shallow and appeared linear. After fitting the gradient with a linear regression, the 1^st^ Max Int gradient had a width of 12.0±3.9 μm. The steeper 2^nd^ Max Int gradient was fit with the logistic function as previously described and had a width of 2.9±1.0 μm. The FWHM gradient across the area scans was found to have a width of 10.5±0.8 μm. There was a significant difference in gradient width between the 2^nd^ Max Int and FWHM gradients as well as the 1^st^ Max Int and 2^nd^ Max Int gradients (alpha=0.05). The location of the 1^st^ Max intensity gradient was shifted from the 2^nd^ Max intensity gradient by 7.6±1.2 µm towards the dentin. The 2^nd^ Max int and the FWHM gradient; however, were centered in the same location with an average shift of only 0.11±0.45 µm.

**Figure 5:**
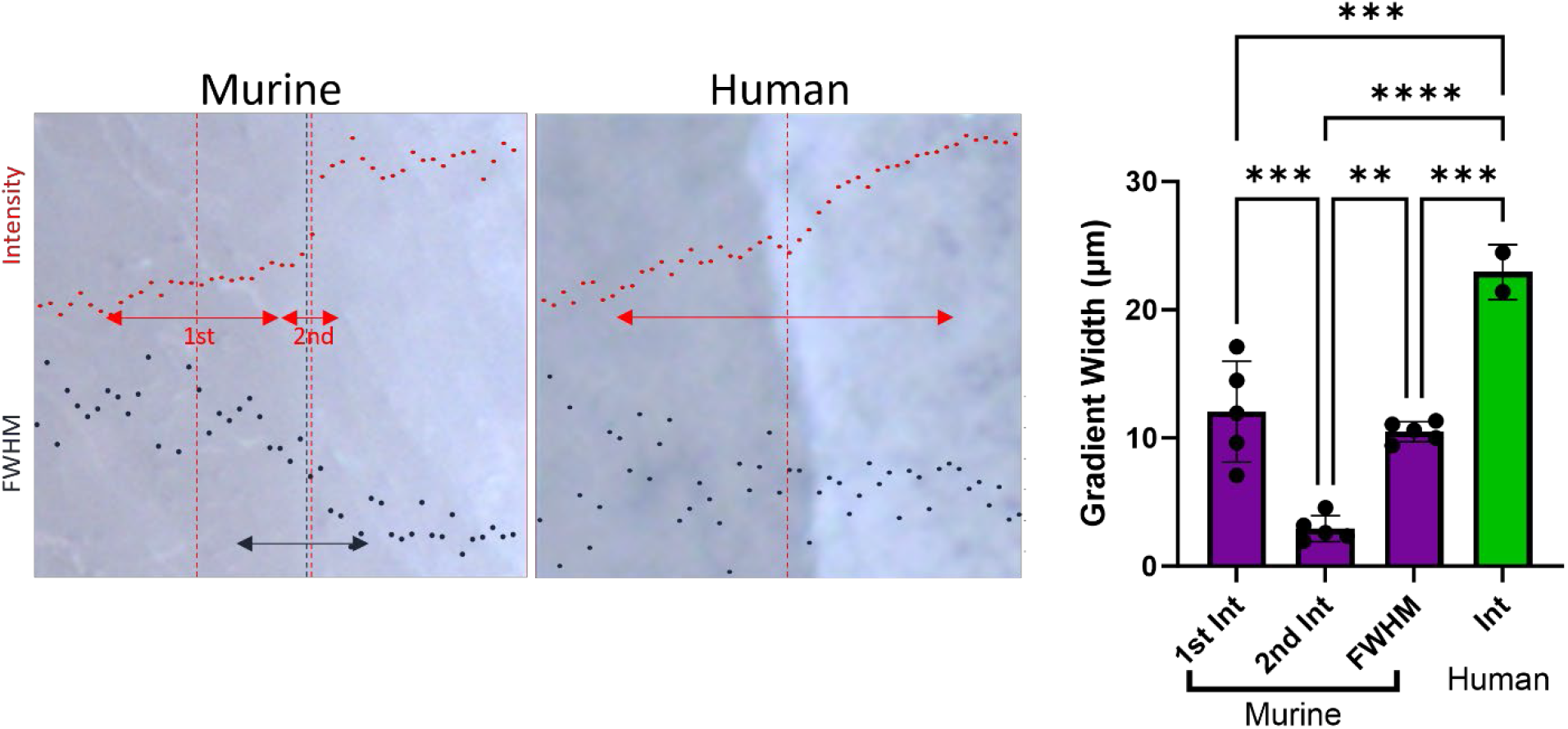
Overlays of representative 960 peak intensity (red) and 960 peak FWHM (black) on birght field images of murine and human DEJs. Arrows represent the width of the measured gradients while the vertical dotted lines represent the center of the respective gradients. (Right) Plot of the measured gradient widths.

The human incisors exhibited a very different gradient geometry, Fig 5. There was a clear gradient in Max peak intensity, but it was much broader than that seen in the murine samples and did not exhibit a two-part gradient. The Max Int gradient width was 24.5±4.2 µm for the first incisor and 21.4±4.3 µm for the second incisor; therefore, the average gradient width for the human incisors was 22.9±2.2 µm. The peak FWHM did not exhibit a measurable gradient across the DEJ and instead remained nearly constant across the 25 µm measured here, although there were greater variations in peak FWHM in the dentin than in the enamel.

### Elastic Modulus of Tissue

For the murine samples, plots of the tissue modulus as a function of position based on the 960-peak intensity as an indicator of mineral content clearly showed that the modulus of the tissue changed as a function of position across the gradient, Fig 6. The starting modulus in the dentin was 57 GPa, which is higher than the values generally accepted for dentin [17], with a final value of 114 GPa in enamel. When the crystallinity was taken into account, the dentin modulus was reduced to 39 GPa while the enamel value remained elevated at 112 GPa. In addition to this change in modulus, the width of the modulus gradient increased when crystallinity was accounted for.

**Figure 6:**
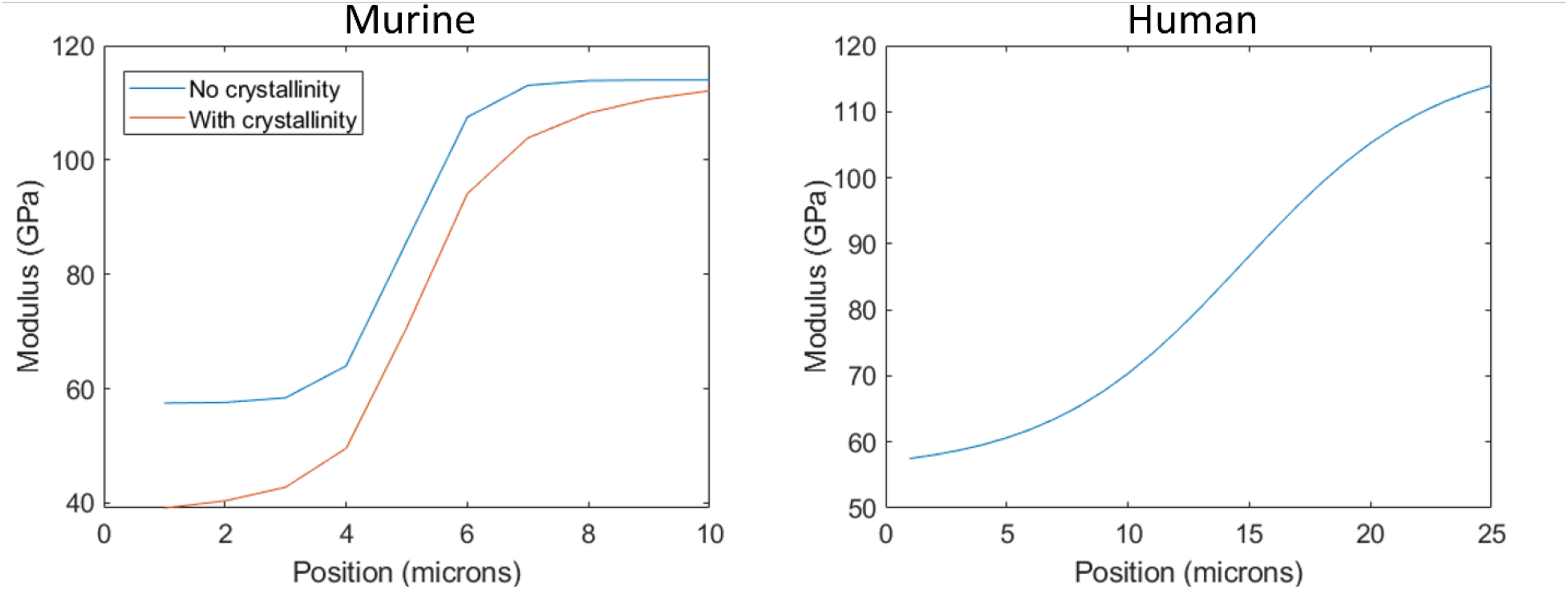
Plots of tissue modulus as a function of position across the DEJ. (Left) Murine DEJ mechanics only considering the mineral content gradient (blue) and considering the crystallinity (red). (Right) Modulus across the human DEJ. Crystallinity is not considered here as there was no change in 960 peak FWHM with position in the human samples.

The average modulus calculated over the width of the DEJ tissue was significantly reduced when the gradient in crystallinity was considered in addition to the gradient in mineral content. It decreased by 13% from an estimated 80.0 GPa to 69.6 GPa.

For the human incisor, there was no gradient in FWHM. Therefore, only the change in mineral content was considered. This results in a modulus value for the dentin of 57.5 GPa and the enamel of 114 GPa in agreement with the set bounds. The average modulus across the 25 µm width was 79.6 GPa.

## Discussion

Human and mouse incisors are both composed of dentin and enamel that meet at the dentin-enamel junction. Despite being composed of the same building blocks, mouse and human incisors exhibit very different structures, growth patterns, and loading regimes. Due to these significant differences in the human and mouse incisors, one would expect the DEJ to exhibit modified structures best adapted to each environment. Here we provide one of the only comparisons of the DEJ structure between murine and human incisors using µCT and Raman spectroscopy and relate it back to possible mechanical benefits.

The width of the DEJ has been reported to be between ∽2-100 µm depending on the location, species, and mode of measurement [3-10]. In this study, two modalities were used to identify different gradients in structure and composition of the DEJ in human and mouse incisors. The first is the use of µCT to measure gradients in density between the species. In this measure, the gradient is determined from the change in attenuation coefficient across the DEJ, which is positively correlated to the density of the tissue. This density gradient was found to be significantly larger in the Human incisors (∽80 µm) than in the murine incisors (∽20 µm), by nearly a factor of four. Due to the resolution of the µCT at 5 and 3.4 µm respectively, this change in density accounts for changes in mineral content but also structural changes such as changes in porosity. Mantle dentin, a thin tissue found between the DEJ and the circumpulpal dentin, is generally shown to be less mineralized than its circumpulpal counterpart [1]. This might point to a density gradient that spans more than the DEJ and instead accounts for the presence of varying thicknesses of mantle dentin between humans and mice. The possibility of including the mantle dentin, sometimes referred to as the sub-DEJ soft zone, in considerations of DEJ structure and mechanics has been previously suggested [4, 18]. Although our Raman results do suggest that there are differences in the width of the non-mantle dentin DEJ, these differences are not large enough to explain the four-fold increase in density gradient width in humans compared to mice. Therefore, we suggest that the wider density gradient in humans is indicative of a wider mantle dentin region than seen in mice.

The second modality used to measure the size of the DEJ in humans and mice was Raman spectroscopy. Prior studies of the DEJ utilizing Raman spectroscopy have reported a human DEJ width of approximately 4-50 µm [19-23], with Xu et al. reporting up to 6 microns of variation in DEJ width within the same tooth when moving from a cervical to occlusal locations [19]. We find that the human incisors exhibit a single gradient in mineral content that spans approximately 24 µm. This is larger than the width reported by some [19-21, 23] but on par with those reported by others [22] for human third molars. A gradient width of 15-25 µm was reported using nano-indentation on human incisors similar to what we see here [6]. However, unlike Slimani et al. who measured a gradient in mineral crystallinity (as determined from the ratio of 960/950 peak areas) across the DEJ, no gradient in crystallinity (as determined from the width of the 960 peak width) was measured for the human incisors in this study [24]. The difference in the ways the crystallinity was determined between the two studies could explain the differences in the results. The 960 peak width is an indicator of atomic order/disorder in the sample while the ratio of the 950/960 peaks provides the relative amounts of “amorphous” to crystalline phosphate in the region of interest. It is possible that the later could exhibit variations while the former does not. This serves as a good reminder of the complexity associated with defining crystallinity.

The murine incisors exhibited a very different structure from the human incisor DEJ. Instead of presenting with a single gradient across the 25×25 µm^2^ area, the mice appear to have a 2-part gradient. The first is a shallow gradient spanning ∽12 µm that has a linear increase in the height of the 960 peak. Following this initial gradual increase in mineral content, there is a rapid rise over the span of ∽3 µm. This second gradient is on the order of certain measures made using high resolution imaging [8, 9]. The sum of the width of the first and second gradients approaches the width of the density gradient measured in µCT. This could suggest that the first gradient, which presents on the dentin side of the DEJ, is indicative of a gradient in mineral content across the mantle dentin while the 2^nd^ gradient is a measure of the mineral content gradient across the DEJ interface. No such two-part gradient was seen in the human incisors; however, that may be due to limited size of the Raman scans (25 µm) as compared to the width of the density gradient (80 µm). Again, unlike the human incisors, the murine DEJ did exhibit a significant change in crystallinity across the DEJ. This gradient in mineral order spanned a width of ∽10 µm and overlapped with the location of the 2^nd^ gradient. Changes in crystallinity across the DEJ have rarely been reported [24] much less interpreted in terms of their contributions to the tissue mechanics.

Recent studies have shown that changes to the mineral composition and mineral crystallinity have significant effects on the mechanical stiffness of the apatite crystals[15, 16, 25]. For example, addition of carbonate substitutions to the apatite lattice significantly reduces crystallinity and in turn reduces the mineral stiffness by up to 50% [15, 16]. Apatite mineral in dentin is known to contain about 3 times the amount of carbonate as apatite in enamel [26] suggesting that this could be responsible for the measured crystallinity gradient in mice. Here, we use that knowledge to determine the relative contributions of gradients in mineral content and mineral crystallinity on the mechanical properties of the DEJ. First, the changes in modulus as a function of position across the DEJ without considering the effect of crystallinity was determined for both the human and murine incisors. Both clearly show an increase in modulus due to increased mineral content from the dentin to the enamel. The width of these gradients is directly proportional to the measured changes in 960 peak height across the DEJ and therefore the width of the modulus gradient is larger in the human samples than in the murine samples. (Note the difference in X-axis between the two plots.) When we average the modulus over the width of the gradient (10 µm for the murine samples and 25 µm for the human samples), we find that the average DEJ modulus is 80.0 GPa for the mice and 79.6 GPa in humans suggesting that despite the differences in structure the overall modulus is preserved. However, when we account for the gradient in crystallinity in the mouse model, we see that the average modulus drops to 69.6 GPa; lower than the values seen in humans. A reduced interfacial modulus is associated with increased crack absorption suggesting that the crystallinity gradient may aid in preventing cracking along the murine DEJ during the high loads associated with gnawing behaviors. Additionally, consideration of the mineral crystallinity widens the mechanical gradient across the DEJ. Wider mechanical gradients are also associated with increased resistance to fracture suggesting that the murine DEJ has developed structures to prevent fracture along this interface. Here we see the importance of crystallinity gradients in controlling the mechanical properties of interfaces between mineralized tissues.

## Conclusions

A combination of Raman spectroscopy and µCT was used to measure compositional and structural gradients across DEJs from mouse and human incisors. Significant differences between the two species were noted including the presence of significantly larger density and mineral content gradients in the humans as compared to mice. Differences in the density gradients as compared to the Raman determined mineral content gradients were determined to be cause by consideration of the mantle dentin in both species. Mice also exhibited significant structural differences compared to humans including the presence of a two-part mineral content gradient and a gradient in mineral crystallinity. Both the gradients in mineral content and crystallinity were used to predict the modulus of the DEJ as a function of position. It was shown that the contributions of changes in crystallinity to the mechanical properties lead to a reduction in the overall tissue modulus as well as a broadening of the mechanical gradient. Both changes in the murine mouse are expected to make the mouse DEJ more resistant to fracture. These results point to the significant differences between human and murine DEJs and the importance of considering both structural and compositional factors when predicting the mechanical behavior of interfacial mineralized tissues such as the DEJ.

